# Utilization of cyanobacterial siderophore cyanochelin B by phylogenetically distant heterotrophs suggest its role in mediating microbial interactions

**DOI:** 10.64898/2026.07.27.740964

**Authors:** Berness Peter Falcao, Martinez Yerena Jose Alberto, Tomáš Galica, Jan Mareš, Clémentine Laffont, Lenka Marešová Štenclová, Surbhi Sharma, Jürgen Tomasch, Divya Aggarwal, Jan Mašek, Petra Divoká, Kateřina Čapková, Tomáš Bešta, Vendula Krynická, Rolf Kümmerli, Pavel Hrouzek

**Author notes:** Correspondence to:* Pavel Hrouzek, Rolf Kümmerli.

## Abstract

Cyanobacteria are key prokaryotic primary producers in diverse ecosystems, yet the role of cyanobacterial siderophores in shaping their associated microbiomes remains unexplored. Our study demonstrates the benefits provided to the heterotrophic co-habitants of filamentous cyanobacteria in terrestrial microbial biofilms, focusing on the recently discovered widespread siderophores cyanochelins.

To address the acceptance of cyanochelin B (CychB) across multiple bacterial classes, we first investigated its role in providing iron to a model siderophore producer *P. aeruginosa* PAO1 and selected *Pseudomonas* natural isolates, which were found to utilize CychB under iron limiting conditions while downregulating endogenous siderophore production. In response to CychB, PAO1 expresses a siderophore internalization cluster, which is localized in multiple *Pseudomonas* natural isolates.

Using metagenome analysis, we characterized the bacterial community recruited along with CychB producing *Phormidesmis* cyanobacteria under long-term iron starvation. Potential CychB acceptor bacteria associated with the CychB producer were predominantly lacking endogenous siderophore machineries. Using siderophore selective pressure, we isolated a genuine CychB acceptor, gram-negative bacterium *Methyloversatilis* sp. S146 and demonstrated that its genome hosts an iron processing cluster overexpressed after CychB feeding, recognizing *Methyloversatilis* as a candidate for further mechanistic investigation of iron acquisition–driven microbial interactions.

Our results indicate that CychB supports a specific subset of co-habiting heterotrophic bacteria during iron starvation, further emphasizing the role of cyanobacteria as key drivers of nutrient flows within globally important microbial soil crust ecosystems, supporting microbial life in nutrient-limited environments. These findings provide a mechanistic foundation to elucidate the role of cyanochelins as a public good in these communities.

## Introduction

Cyanobacteria are important primary producers in aquatic and terrestrial microbial communities, contributing to carbon cycling and maintaining atmospheric oxygen levels [1, 2]. Although they have a high demand for iron, they often inhabit environments where iron is oxidized and precipitates as highly insoluble ferric oxides rendering it “*locked away*” in aerobic condition [3–6]. It is assumed that cyanobacteria, like many heterotrophic bacteria, synthesize siderophores to chelate ferric iron from the environment as principal strategy for iron acquisition [7–9]. However, even though cyanobacteria are widespread in nature and are a reservoir of secondary metabolites, a surprisingly small siderophore repertoire has been reported so far. In stark contrast, more than 1000 chemically different siderophores have been characterized from heterotrophic bacteria.

In natural communities, siderophores can serve as *‘public goods*’ as they are secreted and can be shared among members that possess matching uptake systems. Conversely, siderophores can also serve as *‘public bads*’ by locking iron away from community members with non-matching uptake systems and thereby monopolizing iron for producers [10, 11]. Moreover, instead of producing siderophores themselves many microbes can also exploit (xeno-)siderophores produced by other community members, which is often called *‘siderophore piracy*’ or *’cheating*’. For this purpose, cheaters harbour genes encoding additional TonB-dependent siderophore transporters (TBDTs) dedicated to xenosiderophore uptake [12–14]. However, xenosiderophore use is not always an exploitative act as it can also lead to a mutualistic exchange of secreted goods [15–17].

The involvement of cyanobacteria in these siderophore-mediated interactions and in shaping the microbial communities are largely unknown [9]. Due to their phototrophic way of life, cyanobacteria often experience carbon and energy overflow, and a substantial proportion of this energy gets invested into storage compounds or the synthesis of secondary metabolites [18, 19]. We therefore predict siderophore synthesis to be relatively cheap and consequently the evolutionary pressure to reduce cheating to be weak, thereby favouring the sharing of siderophores as a public good. This differs fundamentally from heterotrophic bacteria, for which siderophore production requires a significant metabolic investment and thus selects for stringent siderophore specificity and cheater control.

To test our hypothesis, we capitalize on our recent discovery of a widely distributed class of siderophores, cyanochelins, frequently featured in genomes of filamentous mat forming cyanobacteria [20, 21]. We have previously chemically characterized and purified cyanobaterial beta-hydroxyaspartete siderophore cyanochelin B (CychB). In our study, we use CychB to test its acceptance by well-characterised *Pseudomonas* strains with well explored siderophore dynamics. Subsequently we have isolated CyChB producer from microbial mat and characterized its phycosphere-associated bacteria occurring under iron starvation which was followed by isolation of CyChB acceptors. The detailed transriptomic study was performed on *Pseudomonas* sp. PAO1 as well as in phycophere-originating bacterium *Methyloversatillis* to further study the effect of CyChB on the acceptor bacterium.

## Results

### CychB is accepted as xenosiderophore by Pseudomonas aeruginosa PAO1

To determine if CychB can be utilized as a xenosiderophore by heterotrophic bacteria, we tested its acceptance by phylogenetically distant *Pseudomonas aeruginosa* PAO1, a model bacterium for both siderophore production and its ability to use xenosiderophores.

In a first experiment, we subjected PAO1 to CychB in the presence of iron (0, 0.5, 0.05, 5, 10, 20µM) and found a negligible decrease in growth integral when CychB was supplemented **(Fig. 1A).** In addition, we quantified the auto-fluorescent pyoverdine, the main siderophore produced by PAO1, but found no detectable production levels **(Fig. 1B**). For iron-rich conditions, we infer that the ferri-CychB complex is likely not imported by PAO1 probably due the overall high iron availability and low siderophore requirements (self- and xeno-siderophores).

**Figure 1.**
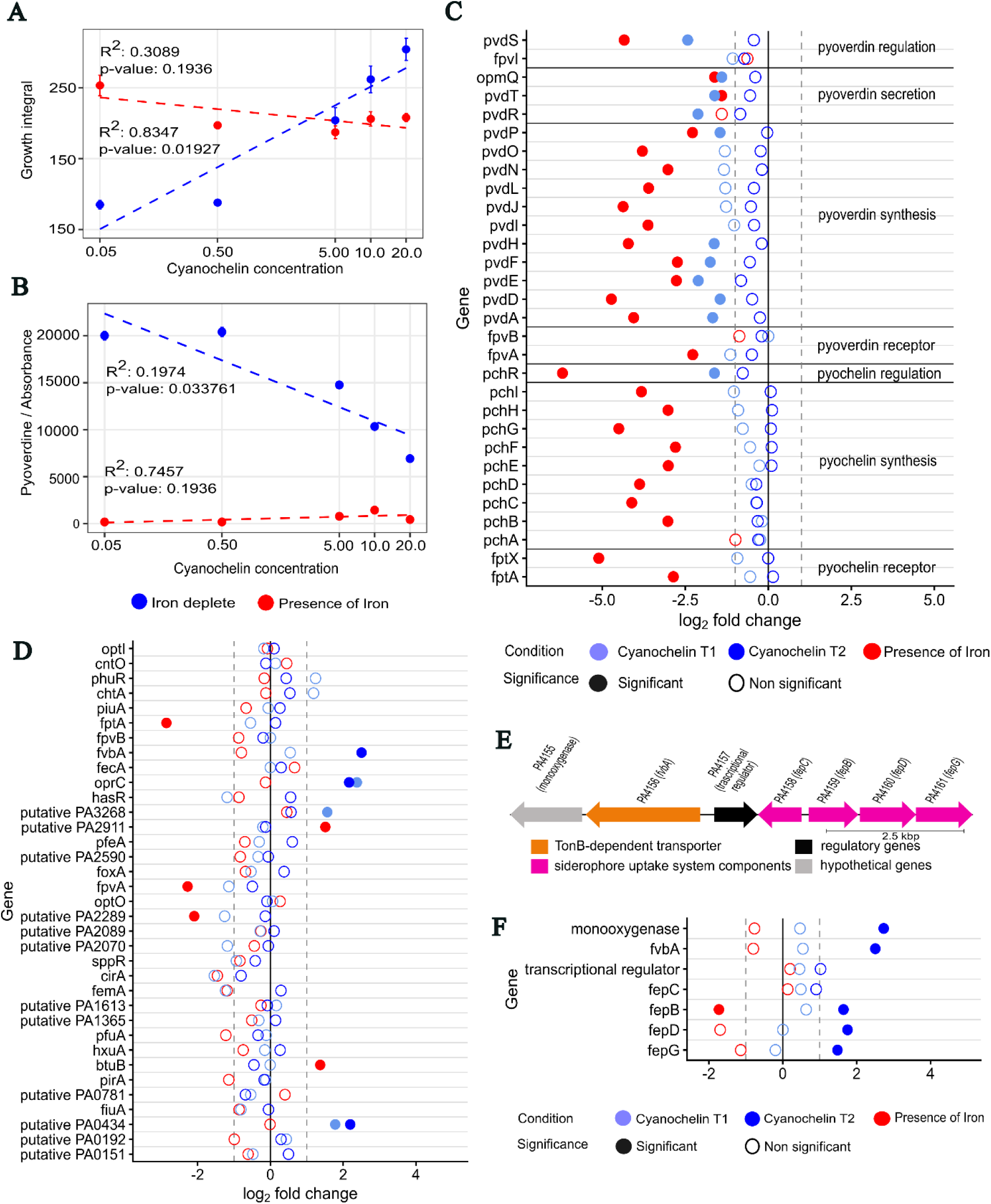
Effect of CychB feeding on *P. aeruginosa* PAO1. **A,** Growth integral for PAO1 culture at different CychB concentration in iron full- and in iron-limited media. Growth curves were measured at OD_600_ over 24 hours and integrated. **B,** Pyoverdine production (area under relative fluorescence unit (RFU) curve divided by the growth integral value) in varying CychB concentration in iron full- and iron-limited media normalized to cell density (OD_600_). Individual data points represent mean values of four replicates. Statistical relationships between the CychB concentration and the measured growth integral/pyoverdine production were evaluated using a linear regression. **C,** Relative expression (log_2_ fold change) of pyoverdine biosynthesis genes at two-time point (T1=6 h, T2=12 h) after CychB treatment. **D,** Relative expression (log_2_ fold change) of all TBDTs present in PAO1 genome after CychB treatment at two-time points (T1=6 h, T2=12 h). **E,** Tentative CychB internalising gene cassette including FvpA (TBDT PA4156). **F,** Relative expression of CychB internalising cassette genes after CychB treatment at T1=6 h, T2=12 h. For C, E, G the full circles represent points that are significant and hollow circles show non-significant values at *p*=0.05.

On the contrary, we recorded a substantial and significant increase in growth integral **(Fig. 1A)** and a decrease in pyoverdine production **(Fig. 1B)** under iron limited conditions with increasing CychB concentration. These results indicate that PAO1 can utilise and benefit from CychB, whilst downscaling the investment in endogenous pyoverdine production.

### RNAseq reveals the upregulation of several TBDTs in response to CychB feeding

To test whether PAO1 expresses specific TBDTs for ferric-CychB import, we conducted a transcriptomic analysis comparing PAO1 gene expression in the presence of iron and in absence of iron along with CychB supplementation. CychB feeding induced a profound transcriptomic response in PAO1 **(Supplementary Data file 1**). When compared to iron starved culture, genes responsible for pyoverdine biosynthesis and transport are downregulated under iron full conditions [22, 23] as well as in iron starved culture supplemented with CychB **(Fig. 1C)**, thus matching our phenotypic data. In addition, we have also recorded downregulation of genes related to pyochelin **(Fig. 1C)**, siderophore produced by PAO1 during moderate iron limitation, which hints strongly that CychB uptake mitigates iron starvation response by endogenous siderophores.

When focusing at TBDTs expression [24], we identified four transporters out of the 35 TBDTs known for PAO1, that were significantly upregulated within 6 (T1) and/or 12 (T2) hours after CychB supplementation **(Fig. 1D)**. OprC precursor (PA3790) was previously shown responsible for importing the cuprous and cupric ions [25, 26]. PA0434 and PA3268 are less well characterized and no other iron internalisation genes are found around these TBDTs. FvbA (PA4156) was previously shown to uptake vibriobactin as xenosiderophore in PAO1 [27]. Notably, this TBDT is located in a strongly upregulated gene cluster along with other iron regulation genes characterized in *E. coli* **(Fig. 1E, F)**. Our results highlight that PAO1 strongly responds to CychB supplementation and reveal four putative CychB internalization pathway in PAO1 **(Supplementary figure 1).**

### CychB also promotes the growth of several environmental Pseudomonas spp

To test whether CychB acceptance and growth promotion is a broader phenomenon, we feed CychB to a panel of 15 natural *Pseudomonas* isolates including both pyoverdine producing (n=11) and non-producing (n=4) strains **(Supplementary Table 1)**. These non-pathogenic *Pseudomonas* isolates originate from soil and freshwater habitats.

In the presence of CychB (20 µM) and under iron starvation, we found that all pyoverdine-producing strains showed increased growth intervals (statistically significant in eight cases) **(Fig. 2A)**. Moreover, seven out of 11 isolates downscaled the production of pyoverdine. Crucially and similar to PAO1, these beneficial effects disappeared under iron-rich conditions (**Fig. 2B**). Although a higher variation in response to CychB is expected among such natural isolates, our results suggest that most of these strains can rapidly recognize the CychB presence, reduce the energetic cost for siderophore production, and switch to the uptake of CychB as xenosiderophore.

**Figure 2.**
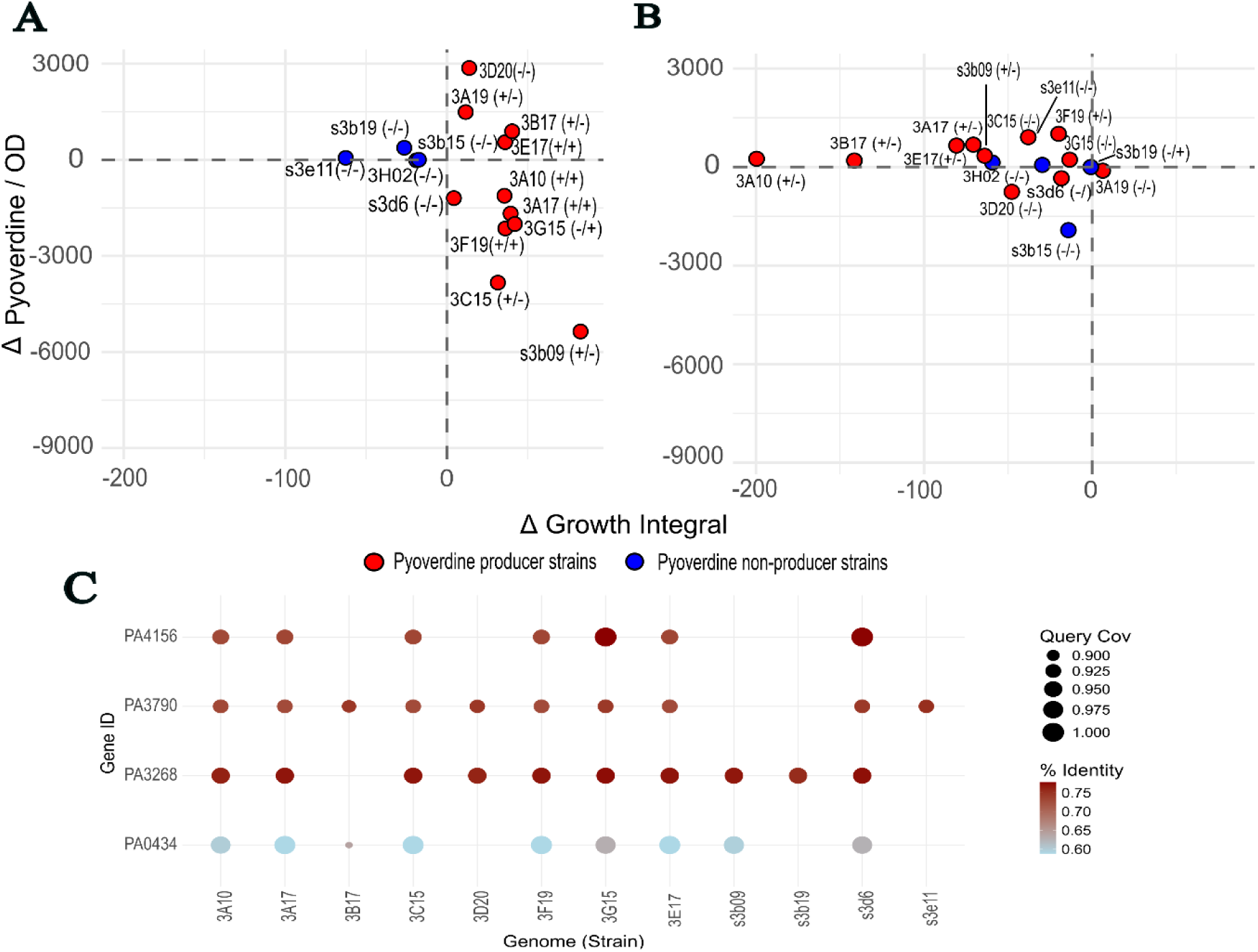
Effect of cyanochelin feeding on *Pseudomonas* natural isolates. Δ Growth Integral and Δ Pyoverdine/OD were calculated as the difference between 20 µM and 0 µM CychB feeding (20 µM − 0 µM) for individual strains in **A,** iron depleted media and **B,** iron full media. Pyoverdine production values were normalized to optical density. Positive values indicate an increase in response under 20 µM CychB relative to control (0 µM). In each of the plots, strains depicted as red circle are pyoverdine producers and strains depicted in blue are pyoverdine non-producers. The symbols in the brackets behind the strain code e.g. (+/-) designates significant difference (pairwise t-test) between the 20 µM CychB and 0 µM CyChB in growth integral (symbol before backslash) and pyoverdine production (symbol after backslash). **C,** Occurrence of PAO1 TBDTs gene homologs upregulated under CychB treatment in *Pseudomonas* natural isolates. Each dot corresponds to a BLAST hit between a query gene (PA4156, PA3790, PA3268 and PA0434) and a *Pseudomonas* genome ORF. Intensity of the dot color reflects percent pairwise identity, and size of the dot represents query coverage. Missing dots show either absence of gene homologs or no hit in the top 10 candidates on BLAST.

In stark contrast, the growth of the four non-pyoverdine producing strains were all reduced in the presence of CychB, suggesting they are not capable of accepting this siderophore **(Fig. 2A)**. These results indicate that the recognition and use of CychB is regulatorily linked to a functional native siderophore system.

Next, we explored whether the natural isolates possess gene homologs of one or several of the four TBDTs found to be upregulated in PAO1 under CychB treatment (**Fig. 2C**). On analysing the top 10 BLAST hits for each gene, the four non-pyoverdine strains had low sequence coverage or lack TBDT homologs. In contrast, homologs to these genes were found in pyoverdine producer strains, suggesting that these strains may possess compatible TBDTs. In addition, to support our hypothesis that TBDT FvbA (PA4156) along with the other genes identified in PAO1 are likely involved in the internalisation of CychB also in other strains, we analysed the *Pseudomonas* natural isolate genomes for homologs of the seven genes in the transport cassette. Four strains (3F19, 3E17, 3A17 and 3C15) showed presence of all 7 upregulated genes involved in the CychB transport cassette identified in PAO1 gene expression. **(Fig1F, Supplementary Fig 2).**

### CychB-producing cyanobacterium Phormidesmis enhances abundance of specific set of cohabitants under iron starvation

Having established the CychB acceptance in *Pseudomonas* model strains, we turned to natural bacterial isolates residing in close proximity to the CychB producer. Specifically, we focussed on cyanobacteria-dominated microbial mats and investigated whether CychB producers confer CychB-mediated iron supply benefits to co-habiting heterotrophic bacteria. For this, we conducted a field sampling to collect cyanobacteria rich mat samples and further subjected them to controlled iron starvation to obtain a CychB positive microbial mat and subsequently isolated the CychB producing cyanobacterium *Phormidesmis sp.* [21].

One fraction of the original cyanobacterial mat sample was preserved in glycerol stock and the other fraction was subjected to iron depletion in Z-medium. The cyanobacterium was isolated from iron depleted culture after confirmation of *CychB* production via single filament picking. The procedure included several washes in order to maintain the microbial community closely associated with *Phormidesmis* filaments. Subsequently the stabilized culture was transferred to Z-medium and iron depleted Z-medium. The original mat sample from field, community culture maintained in iron limited condition, and *Phormidesmis* culture in Z-medium with/without iron underwent metagenomic analysis to identify the microbial community. As expected, the microbial community of the original microbial mat is significantly more diverse than all the obtained lab cultures **(Fig 3A, B)**.

**Figure 3.**
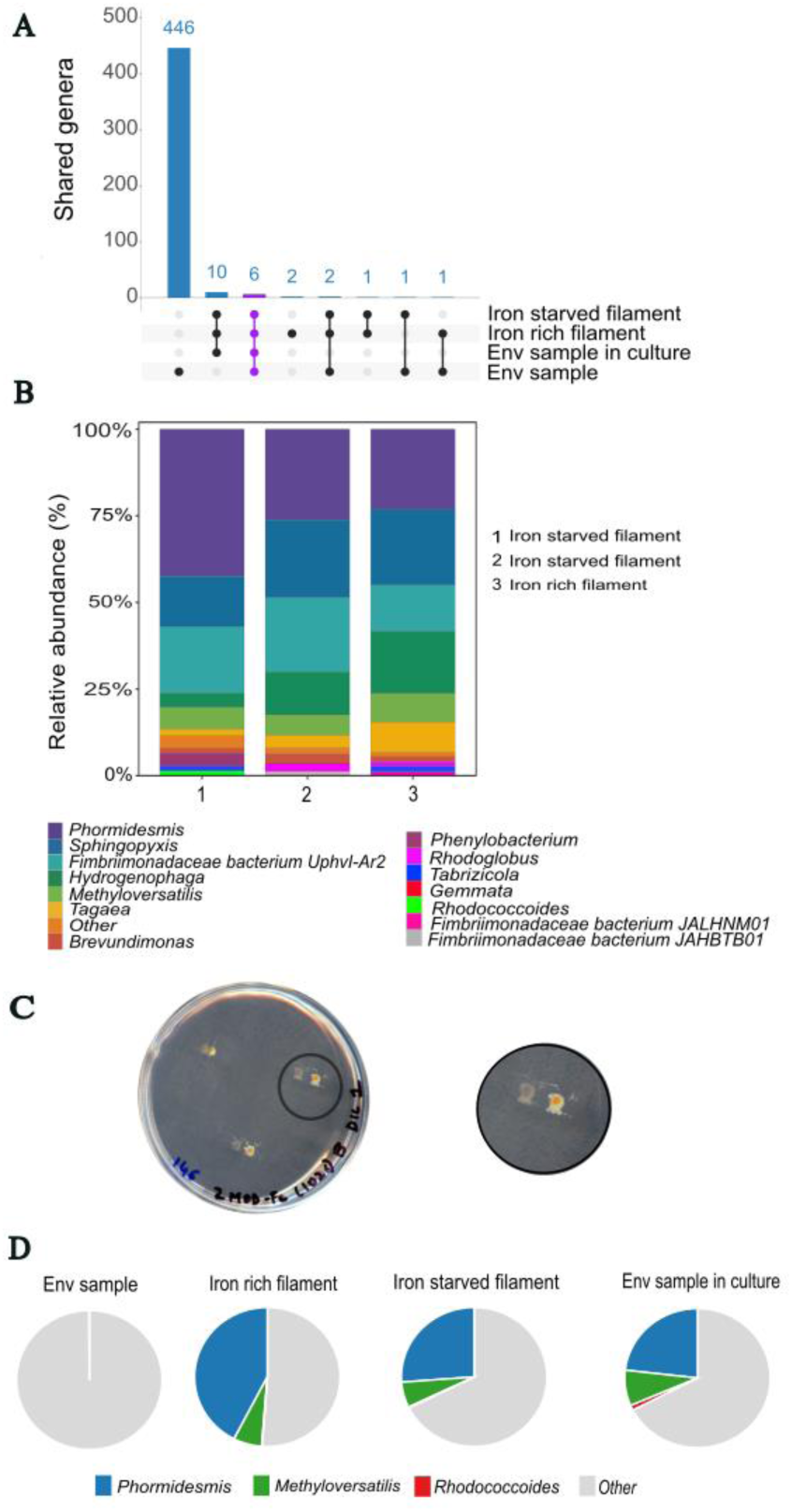
Metagenomic analysis of collected cyanobacterial mat (Env sample), sample of cyanobacterial mat long-term cultured in iron starved conditions (Env sample in culture) and cyanochelin producer (*Phormidesmis*) culture in Fe full conditions (Iron-rich filament) and iron-starved conditions (Iron-starved filament). **A,** UpSet plot showing number of unique or shared genera between samples. Vertical bars represent number of genera and nodes/lines below the graphs show number of genera in individual samples or shared between the samples. **B,** Stacked bar plot showing the relative abundance of the top 10 genera (per sample) in Iron-rich filament, Iron-starved filament, and Env sample in culture. Relative abundance is calculated as the proportion of reads of individual genus to total number or reads within particular sample. **C,** Isolation of *Methyloversatilis sp.* S146-33 and *Rhodococcoides sp.* S146-33 on the Petri dish containing Z modified medium supplements with 20uM cyanochelin. The bacteria growing in the vicinity of the iron bead were considered as CychB acceptors. **D,** Pie charts showing presence/abundance of the CychB producer (*Phormidesmis*) and the two cohabitants (*Methyloversatilis sp.* S146-33 and *Rhodococcoides sp.* S146-33).

We have analysed the overlapping taxa among original microbial mat and *Phormidesmis* filament cultures. Six genera, *Phormidesmis, Sphingopyxis, Hydrogenophaga, Tabrizicola, Brevundimonas,* and *Fimbriimonas* were present in the original environmental sample of the microbial mat, the iron depleted environmental sample culture, and the laboratory *Phormidesmis* filament cultures with or without iron **(Fig. 3 A,B)**. We recovered metagenome-assembled genomes (MAGs) of these taxa and analyzed their genomes for the presence of biosynthetic gene clusters (BGCs) of metallophores. Notably only the *Hydrogenophaga* genome contained a BGC for synthesis of an endogenous enterobactin-like siderophore **(Supplementary Fig 3)**. This points to a significant proportion of bacteria associated in the *Phormidesmis* phycosphere lacking siderophores. Further, Methyloversatilis, *Tagaea*, *Fimbriimonadaceae* bacterium (JAHBTB01, JALHNM0, JALOLT01, uphvi-ar2), *Mesorhizobium* sp. RCIZ01, *Chthonomonadaceae* bacterium JAEUTO01, *Rhodococcoides* and *Rhodoglobus* were detected in *Phormidesmis* laboratory cultures but not in the original metagenome. These bacteria were either enriched with the available media components or could be contaminants occurring during laboratory sample processing and likely benefit from *Phormidesmis* exudates. Again, out of 10 inspected metagenomes only *Rhodococcoides* sp. MAG contained the BGC for an unknown phenolate-hydroxamate siderophore (**Supplementary Fig 3**). Notably in case of *Mesorhizobium*, *Tagea*, and *Hydrogenophaga*, their iron-processing clusters contained TBDT transporters along with *hmu* TUV genes **(Supplementary Fig 3)** analogous to that detected in *Methyloversatilis* (see later).

To isolate cohabitants benefiting from CychB, we applied siderophore selective pressure in combination with presence of a localized source of immobilized iron (FeCl_3_ immobilized in alginate beads) to create a zone of siderophore-iron complex that enriches the growth of cohabiting CychB-benefiting bacteria **(Fig. 3C)**. Using this approach, we successfully obtained gram-positive bacterium *Rhodococcoides sp*.

S146-33 and gram-negative bacterium *Methyloversatilis sp. S146-33* benefiting from CychB. While *Rhodococcoides sp.* S146-33 formed only a minor constituent, *Methyloversatilis sp.* S146-33 was substantially more enriched in *Phormidesmis* lab cultures **(Fig. 3D)**. Since the metagenomic data did not detect these bacteria in the original mat, we designed specific PCR primers and confirmed the presence of *Methyloversatilis sp.* S146-33 but not *Rhodococcoides sp.* S146-33 in the original community **(Supplementary Figure 4).**

### Rhodococcoides sp. and Methyloversatilis sp., cohabitants of CychB producer, gain growth benefits in presence of CychB

In order to confirm that *Rhodococcoides sp. S146-33 and Methyloversatilis sp.* S146-33 can procure CychB-mediated iron under starvation. We first carried out CychB feeding experiments, followed by a genome analysis of both the strains. As *Methyloversatilis sp.* S146-33 was identified as a naturally occurring cohabitant, we primarily focused on characterization of this strain. Isolated strain formed flakes in liquid medium which hampered the direct OD values measurement. It is a common trait observed in *Methyloversatilis* strains EHg5, FAM5T and 500 to clump together in liquid medium and form white flakes [28]. Therefore, we performed the experiments using solid media. An apparent growth was recorded on agarose Z-medium plates with presence of 2.5µM CychB, while *Methyloversatilis* did not proliferate well on iron depleted plates (**Fig. 4A**). This hints that the presence of CychB positively affects the growth of *Methyloversatilis*.

**Figure 4.**
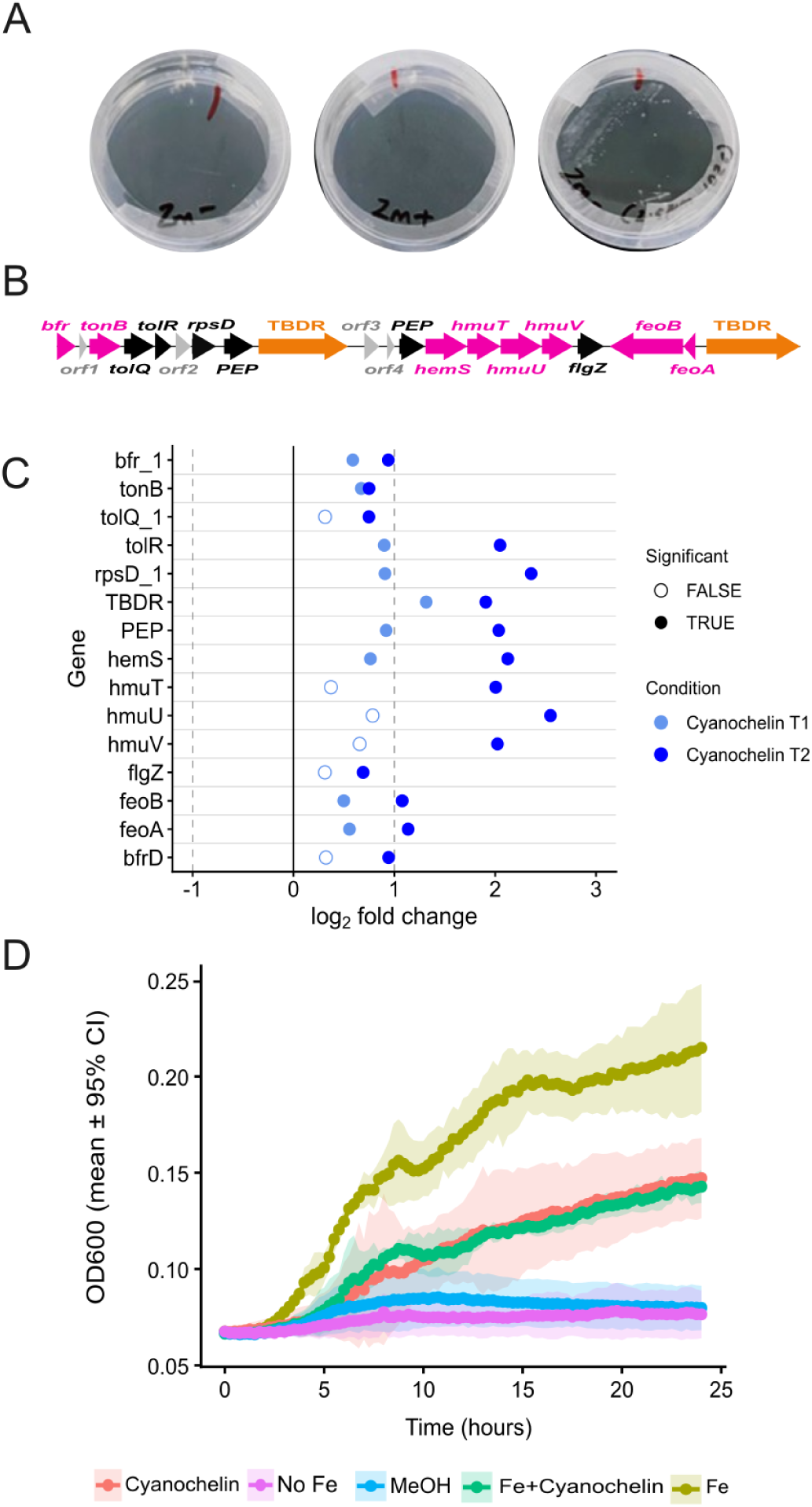
*Methyloversatilis sp.* S146-33 and *Rhodococcoides sp.* S146-33 accept cyanochelin B. **A,** *Methyloversatilis sp.* S146-33 plated on agarose Z-medium plates iron depleted plates and iron replete plates and in presence of 2.5µM CychB. **B,** Iron processing cluster present in *Methyloversatilis sp.* S146-33 strain and its closest relative *Methyloversatilis discipulorum* showing and array of genes for iron import. **C,** Relative expression of iron processing cluster after CychB treatment at T1=8h (light blue) and T2=24 h (dark blue). Each dot represents the log2FC of a gene; y axis showing locus tags of the genes. Full circles represent points that are significant and hollow circles show non-significant values at *p*=0.05. **D,** *Rhodococcoides sp.* S146-33 growth curves of cultures grown with/without Fe/ CychB measured as OD_600_ over 24 h. Each line represents a different condition and shaded areas represent 95% confidence intervals.

Further, we analyzed the genomes for possible TBDT transporters involved in CychB import as well as to characterize the effect of pure CychB on the bacterium. We identified a gene cluster containing two TBDT transporters colocalized along with set of iron transport genes including heme/hemin transporter genes (hmuTUV), tonB, FeoB (responsible for Fe2+ uptake), and bacterioferritin (**Fig. 4B**). To map the response of *Methyloversatilis* to CychB, we performed a transcriptomic experiment. In total **77** genes were significantly altered after 24 h treatment **(Supplementary Data file 2).** As expected, the full iron processing cluster was significantly upregulated after CychB treatment, supporting its putative role in CychB import (**Fig. 4C**).

To determine if CychB had a positive effect on *Rhodococcoides* growth, we performed a feeding experiment and monitoring of cell growth for 24 hours. As expected *Rhodococcoides* is capable of proliferating in the presence of insoluble iron source (FeCl_3_) in the media. However, under iron starved conditions negligible growth was recorded **(Fig. 4D)**. Supplementation with CychB clearly recovered its growth even in iron starved conditions to a similar degree as observed in *Pseudomonas* strains, reaching approximately half of the growth rate compared to iron starved conditions. The analysis of the *Rhodococcoides* genome identified the biosynthetic gene cluster for hydoxamate-salicylate type of siderophore (**Supplementary Figure 3**). However, our supplementation experiments clearly show that under short-term iron limiting conditions the bacterium is not able to restore its growth using its own siderophore machinery and benefits from externally supplemented CychB.

In conclusion, our experiments on the isolated cohabitants *Rhodococcoides* sp. S146-33 and *Methyloversatilis* sp. S146-33 prove that *Phormidesmis* cohabitants are capable of processing CychB to support their growth and survival. This is further supported by the lack of siderophore biosynthetic machinery in many of the bacteria in the phycosphere of this cyanobacterium, which may indicate to dependence on CychB as a xenosiderophore

## Discussion

Heterotrophic bacteria are involved in diverse microbial interactions like cooperation, exploitation and competition via specific siderophore–transporter pairs, and optimize their iron-acquisition strategies within the complex community by creating a dense metabolic network connecting different species, genera, families or even kingdoms [12, 29]. Despite the essential role of cyanobacteria as primary producers and keystone species in many microbial communities, cyanobacterial siderophores and cyanobacterial-siderophore mediated biological interaction have gained little attention [30–32]. It is reasonable to expect that cyanobacteria are potential candidates for siderophore sharing, as they have easy access to ATP and reductive equivalent supplied from photosynthesis, that could support siderophore biosynthesis and its distribution to other community members [18].

Our study focused on biological interaction mediated by CychB, a representative of cyanobacterial beta-hydroxyaspartate containing siderophores [20, 21]. We provided evidence that CychB is accepted as xenosiderophore by a range of phylogenetically distant heterotrophic bacteria including gram negative *Pseudomonas* spp. and *Methyloversatilis* sp. S146-33 strains as well as a gram-positive bacterium *Rhodococcoides* sp. S146-33.

We initially used *P. aeruginosa* PAO1 as a model strain to study xenosiderophore uptake. PAO1 is known to uptake structurally diverse xenosiderophore such as mycobactins, enterobactin, schizokinen, using multiple well-characterized TonB-dependent transporters [33]. From previous studies we also have evidence that different *Pseudomonas* strains react differently to the presence of a xenosiderophore based on the available TBDT set. Some *P. aeruginosa* strains can use ferrichrome and enterobactin, but cannot utilize structurally similar bacillibactin [29, 34, 35].

In our study, uptake of CychB by PAO1 is likely via the FvbA-containing cluster overexpressed under CychB treatment. Notably, this cluster was also conserved in 4 of the natural isolates that increased growth and reduced pyoverdine production in presence of CychB. Interestingly the non-pyoverdine producers did not gain any benefits from CychB. Thus, more efforts need to be taken to understand the interlink between the pyoverdine production and genes for xenosiderophore uptake in Pseudomonas. Pseudomonads are found within a microbial community of cyanobacteria dominated soil crusts in deserts. Even though low abundances were recorded, they are estimated to functionally drive iron acquisition for the community [36]. Reduction of pyoverdine production in presence of CychB, as seen in our study, opens a question whether *Pseudomonas* act like donors or acceptors of siderophores in a cyanobacterial rich community.

We previously demonstrated that CychB biosynthetic gene clusters are frequently found in filamentous cyanobacteria belonging to the *Leptolyngbyaceae* family, inhabiting microbial mats and soil crusts [20]. Since these potential CychB donors are major primary producers found in soil habitats of deserts and steppes, the importance of broad acceptance of CychB as xenosiderophore by distinct heterotrophs in the community provides the basis for further investigation of their ecological importance. It is worth mentioning that CychB-like BGCs were found also in other filamentous cyanobacterial species, including *Microcoleus vaginatus*, the key ecosystem engineer species in biological crusts of arid and semi-arid environments across the globe [21, 37, 38]

Because of this potentially wide impact, we focused on investigation of heterotrophic epiphytic microzone of a CychB producer acquired from the field. Our data corroborates that *Phormidesmis* culture enriched under iron starvation possess phycosphere associated bacteria benefiting from CychB production under iron starvation as exemplified in case of the genuine natural CychB acceptor *Methyloversatilis*. The closest relative of the isolated strain *Methyloversatilis* sp. S146-33 is *Methyloversatilis discipulorum* originated from the sediment of the freshwater Lake Washington. However, the global distribution of closely related 16S rRNA sequences showcase that *Methyloversatilis* can be found in wet soils, wastewater biofilms, lake sediments, plant or animal related microhabitats [39]. Notably, the *Methyloversatilis discipulorum* genome lacks BGC for siderophore production and possesses the same iron processing cluster found in S146-33.

The fact that many of the cohabitants, including *Methyloversatilis*, do not possess siderophore biosynthetic machinery suggests that these bacteria rely on iron sources provided in bioavailable form by other organisms, likely CychB in this case. Although it is difficult to reconstruct the exact evolutionary trajectories which might involve adaptive loss of siderophore BGC or lateral gene transfer of siderophore transporters, it is plausible to presume that *Phormidesmis* cohabitants evolved under stable siderophore income. For instance, experimental studies on *Pseudomonas* demonstrated rapid loss of siderophore BGC leading to clones that cheat on clones still producing pyoverdine as the cheater still maintains compatible transporters [40]. In extreme situations, acceptance of xenosiderophore can lead to a direct dependency of siderophore non-producing bacteria dependent on siderophores produced by other bacteria [41].

It is worth mentioning that CychB, like all known beta-hydroxy aspartate siderophores, undergo photolytic cleavage in presence of UV, catalysing the reduction of Fe^3+^ to biologically available Fe^2+^. Previously we have demonstrated that the CychB producer *Leptolyngbya* sp. NIES-3755 supports the growth of the model cyanobacterium *Synechocystis* sp. PCC 6803, which lacks siderophores, in presence of UV light in vitro [21]. This phenomenon adds another layer of complexity to the microbe-microbe interaction as the photolytically reduced iron is readily available to all members of the community living in the close vicinity of *Phormidesmis* filament. This applies for *Phormidesmis* co-habitants, lacking endogenous siderophores, obtained from the culture derived from a single cyanobacterial filament after multiple washes indicating close association of the heterotrophic bacteria with the cyanobacterium. In our manipulative experiment, we have on purpose studied acceptance of the intact molecule of CychB, i.e. under total absence of the UV lytic light. Irrespective of its photolytic nature, the existence of dedicated transport machineries in *Pseudomonas* and *Methyloversatilis* demonstrates the benefits of intact CychB. The acceptance of non-photolyzed CychB by other heterotrophic bacteria in nature might bring advantage during the dark periods of days or in deeper zones of the cyanobacterial soil crusts rich in UV-protective compounds. However, we expect both intact CychB as well as photolytically reduced iron to favour the bacterial cohabitants of CychB producers.

Interestingly, heterotrophic cohabitants (*Mesorhizobium*, *Tagea*, and *Hydrogenophaga*) including *Methyloversatilis* that reside with the *Phormidesmis* filament show the presence of hemin uptake genes (*hmuTUV*) and TBDT clustered in the genome. These genes have been reported to import heme bound iron from decaying plant material in Rhizobia species [42]. Lysis of cyanobacteria in rich crusts was shown to release intracellular hemoproteins such as cytochromes, thereby providing a source of heme in the environment [43]. This indicates that Phormidesmis cohabitants follow multiple strategies to gain iron during iron starvation alongside the intact CychB acceptance or photolysis of the compound.

To conclude, we have convincingly demonstrated that CychB are utilized as a public good for diverse heterotrophic bacteria. It is reasonable to presume that cyanochelins can be important players in microbe-microbe interactions within globally important microbial mats and crust ecosystem, where the majority of so far known cyanochelin producers resides. In the case of phycosphere associated bacteria the future research should be directed towards establishing whether this siderophore exchange is a part of broader mutualistic relationship between heterotrophic bacteria and cyanobacteria.

## Material and methods

### Determining the benefits of CychB feeding on Pseudomonas natural isolates

*Pseudomonas aeruginosa* (PAO1) and *Pseudomonas* natural isolates from Rolf Kummerli’s laboratory, University of Zurich were used in this study. Genomic data of natural *Pseudomonas* isolates were obtained from an established collection of 315 pseudomonads, isolated from eight soil and eight freshwater (pond) collections spots (18–20 isolates per sample) from the Irchel Campus, University of Zurich.

The selected *Pseudomonas* strains (15 natural isolates) were revived from glycerol stocks and cultured overnight in Lysogeny Broth (LB) at 30°C with shaking at 200 rpm. Overnight cultures were centrifuged at 5,000 × g for 5 minutes and washed with sterile LB medium. The washed cells were resuspended in iron-free BG11 and adjusted to a final OD600 of 0.005 for inoculation. BG11 media supplemented with FeCl3 (∼20 µM) and 25 mM HEPES buffer and BG11 medium lacking added iron, amended with 250 µM 2,2’-bipyridyl (iron chelator) and 25 mM HEPES buffer were used to test the impact of CychB supplementation. CychB was added to cultures at final concentrations of 20 µM. For each condition, 200 µL of inoculated medium was dispensed into wells of a sterile 96-well plate. Plates were incubated at 28°C in Tecan and growth curves were calculated with OD600 readouts obtained every 15 minutes for 24 hours and raw pyoverdine production values were simultaneously obtained at an excitation wavelength of 400nm and emission wavelength of 460nm.

### Metagenome and whole genome sequencing and bioinformatic analysis of CychB-producing samples, cultures, and Phormidesmis cohabitants

Metagenomic data was obtained at 3 consecutive stages of cyanobacterial strain cultivation: (A) The sample collected directly from the field, (B) The culture maintained in the culture collection from the field in iron starved condition, and monoclonal cultures established from single filaments of the CychB-producing *Phormidesmis* sp. S146-33 in iron starved conditions (C) and iron rich conditions (D). Total metagenomic DNA was extracted using the NucleoSpin Soil Mini Kit (Macherey-Nagel, Düren, Germany) with lysis buffer S2 and SX enhancer in duplicates and pooled to reduce sample heterogeneity bias. mDNA integrity, purity, and concentration were determined by UV-gel electrophoresis (TAE buffer, 70V, 1 h), Qubit 3.0 Fluorometer (Thermo Fisher Sceintific, Waltham, USA) broad-range dsDNA assay, and spectrophotometric measurement (BioSpec-nano - Shimadzu, Duisburg, Germany). Sequencing was performed by the Institute of Biophysics, Czech Academy of Sciences using the Illumina NovaSeq X Plus platform (Illumina, San Diego, CA, USA) and pair-end libraries, the sequencing yields of 50 Gbp (150 bp PE library) and 10 Gbp (300 bp PE library) were used for the original biofilm community and the laboratory cultures, respectively. The metagenomes were quality-filtered, assembled, and binned using the standardized *nf-core/mag* v. 4.0.0 pipeline [44] with MEGAHIT v. 1.2.9 assembly algorithm [45] and MaxBin2 v. 2.2.7 binning procedure [46], taxonomy of contigs and bins was assigned by GTDB-tk version 2.4.0 [47]. The content of NRPS/PKS-encoded metallophore BGCs was determined using antiSMASH v. 8 [48] and further annotated using MIBiG 4.0 [49] and custom BLASTp/CDD analyses [50] to assign biosynthetic and transporter protein functions. Taxonomic profiling analysis was performed using the SingleM pipeline v. 0.21.3 [51] yielding OTU tables with relative abundance of taxa hierarchically classified according to GTDB. Taxa abundances and coincidences across samples were visualized using R 4.4.1 [52].

Bacteria isolated from *Phormidesmis* sp. S146-33 iron-starved cultures (*Methyloversatilis* sp. S146-33 and *Rhodococcoides* sp. S146-33) were grown in batch culture and harvested for gDNA isolation by centrifugation. The DNA isolation and genome sequencing procedures followed those described earlier in the text for metagenome sequencing, but the sequencing depth was adjusted to single bacterial genomes (around 5 Gbp, 300 bp PE library). Whole genomes were assembled using the *nf-core/mag* pipeline as above but using SPAdes v. 4.1.0 [53] for contig assembly; genomic contigs were analyzed using antiSMASH v. 8 and custom BLASTp analyses to assess the content metallophore BGCs and TonB-dependent siderophore transport systems as previously [21, 54]. BGCs and siderophore transporter gene cassettes were mapped and visualized using clinker v. 0.0.32 [55].

The metagenome sequencing results were deposited and are publicly available through the European Nucleotide Archive (ENA), study accession PRJEB120791, containing the raw sequencing read archives (ERR17544105, ERR17543440, ERR17543932, and ERR17544083), primary metagenomic assemblies (ERZ29732119, ERZ29732417-419), binned assemblies (ERZ29732429-721, ERZ29732752-776, ERZ29732782-792), and MAG assemblies (GCA_986917915, GCA_986917985, GCA_986917725, GCA_986917735, GCA_986917705, GCA_986917715, GCA_986917975, GCA_986918505, GCA_986917845, GCA_986918035, GCA_986918575, GCA_986917815). Whole genome sequences of isolated *Phormidesmis* cohabitants were deposited through NCBI GenBank as *Rhodococcoides* sp. S146-33 (JCAJWY000000000) and *Methyloversatilis* sp. S146-33 (JCANOT000000000).

### Isolation of CychB producers and cohabiting CychB-benefiting bacteria

Once the Phormidesmis filament enriched culture with confirmed CychB production was obtained, it was used to isolate the cohabiting bacteria. Culture was homogenized using ultra-sound and filtered using a 20 µm pluriStrainer (pluriSelect, Leipzig, Germany). The culture in suspension was diluted 100 and 1000 times and plated on a petri dish with Z modified medium **(Supplementary Table 2)**. This media was supplemented with 20uM CychB; a precipitated FeCl_3_ immobilized alginate bead [21] was placed on the agarose surface to the centre of the plate. The colonies of bacteria growing explicitly around the bead were collected and serially diluted to obtain a single colony of the bacteria present followed by verification of the strain quality using 16S PCR assessment. Whole genome sequencing was performed to determine the identity of the bacteria. The axenic bacteria were preserved in 25% glycerol until further use.

### Analyzing the genomes of Phormidesmis sp. S146-33 cohabitants: Rhodococcoides *and* Methyloversatilis

A tBLASTn search was conducted using all the transporting proteins on the plasmid containing the complete biosynthetic gene cluster for CychB on the Leptolyngbya sp. NIES-3755 strain (AP017310.1). The results were manually inspected to identify clustered hits for the queried genes. The largest contiguous region containing hits for the CychB transporting genes were blasted against the NCBI database to find closely related taxa containing the queried region. As mentioned above, like for *Methyloversatilis* genomes, presence of any siderophore biosynthetic gene clusters in *Rhodococcoides* was identified using the antiSMASH web server v. 8 [48] with default settings, and clusters annotated as NRPS/NRPS-like metallophore or siderophore were recorded.

### Confirmation of the origin of Methyloversatilis sp. S146-33

The identity of *Methyloversatilis sp.* S146-33 as a true cohabitant of *Phormidesmis sp.* S146-33 was confirmed by a two-step nested PCR approach designed to selectively amplify a fragment of a TonB-dependent receptor gene from the clustered region mentioned above. Primer pairs for the target gene were designed on Geneious Prime 2022.1.1 using the our *Methyloversatilis* sp. S146-33 genome as template (NCBI accession JCANOT000000000).). All PCR reactions were performed using PPP Master Mix (Top-Bio, Czech Republic; Cat. No. P125) with 0.25 µM of each primer, and were consequently purified using the NucleoSpin Gel and PCR Clean-up kit (Macherey–Nagel, Germany; Cat. No. 740609.50)

The first PCR reaction contained 1 µg of DNA extracted from the environmental sample from which both *Phormidesmis* sp. S146-33 and *Methyloversatilis* sp. S146-33 were isolated, and were performed using primers jumpF (CTTCCACGTCGAACAGATC) and jumpR (CGACTCCTACAGCTTCTAC), amplifying a 919 bp fragment of the target gene. Amplification was carried out using a touchdown PCR protocol consisting of an initial denaturation at 95°C for 5 min, followed by 10 cycles of denaturation (95°C for 30 s), annealing (30 s, decreasing from 65°C to 58°C), and extension (72°C for 1 min), followed by 20 cycles with a constant annealing temperature of 58°C. A final extension step was performed at 72°C for 5 min.

The second PCR reactions contained 0.1 µg of purified first-round amplicon as template, and was performed using primers MV-TBDT-F (CATCTTTACTACACGACGAAGA) and MV-TBDT-R (AACAGAACGACACCAACAT) to amplify a 167 bp region within the first PCR product. The second PCR consisted of an initial denaturation at 95°C for 2 min, followed by 35 cycles of denaturation (95°C for 30 s), annealing (58°C for 30 s), and extension (72°C for 30 s), with a final extension at 72°C for 2 min. Finally, the sequence of the final amplicons was confirmed by Sanger sequencing and aligned against its corresponding TonB-dependent receptor sequence at the *Methyloversatilis* sp. S146-33 genome to confirm the origin of the strain.

### Determining the benefits of CychB feeding on the isolated bacteria (cohabitants of the CychB producer)

*Rhodococcoides* sp. S146-33 was revived from glycerol stocks and grown overnight in LB medium at 30°C shaker conditions. Culture was diluted in 0.5% CAA medium to a starting OD600 of 0.005 and incubated under different conditions in BG-modified medium. In the test, 4uM of the compound was added to the growing culture. Culture was also subjected to CychB: iron complexes (2:1 ratio). In control, culture was maintained in 4uM iron and methanol (% of methanol in CychB). Growth kinetics of the cultures were monitored for 24 hours with absorbance measurements taken every 15 minutes. Optical density readings at 600 nm were recorded using a *FLUOstar*^®^ *Omega* multi-mode microplate reader. The Growth curve and growth integrals plots were generated using ggplot2 in R 4.4.1 [52, 56]

### Differential expression analysis of Methyloversatilis and PAO1 in the presence and absence of Xenosiderophore

*Methyloversatilis* cultures were initially grown in Min-E medium [57] at 25°C for 48 h. Cells were subsequently transferred to CAA medium supplemented either with iron or without iron. Iron-depleted cultures were supplemented with 0.8% (w/v) NaCl and adjusted to an OD_600_ 0.1. Where indicated, cultures were treated with CychB at a final concentration of 20 μM, while untreated cultures served as controls. Samples of *Methyloversatilis* were collected at the following time points: immediately after transfer to CAA medium containing iron (T0, iron-replete control); immediately after transfer to iron-depleted CAA medium (T0, iron-depleted control); and following incubation under iron-depleted conditions for 8 h (T1) and 24 h (T2) in the presence or absence of CychB.

For *Pseudomonas aeruginosa* PAO1, cultures were grown in LB medium for 24 h before transfer to CAA medium for an additional 24 h. Experimental treatments were performed as described for *Methyloversatilis*. Samples were collected immediately following transfer to CAA medium (T0), and after 6 h (T1) and 12 h (T2) of incubation.

For total RNA, 5 ml aliquot of *Methyloversatilis* cells was pelleted by centrifugation at 5000 rpm for 2 minutes. The resulting cell pellets were immediately processed for RNA extraction by bead beating in PGTX. Cells were broken mechanically in a bead beater using silica beads (zirconia: 100-200 μm) in PGTX with lysis cycle of 3 x 20 s with a 2 min pause on ice between cycles. However, for PAO1, 5 ml aliquot was vacuum-filtered onto hydrophilic polyethersulfone filters (Supor 800, 0.8 µm), and the filters were immediately immersed in 0.8 ml PGTX [58, 59] and vortexed for 10-15 s, followed by an incubation at 95°C for 5 min and chilling on ice for 10 min.

All samples were then extracted twice with equal volumes of chloroform, each followed by centrifugation at 3000 x g for 3 min at RT. The uppermost phase was mixed with an equal volume of isopropanol and incubated overnight at -80°C. RNA was pelleted by centrifugation at 12000 x g for 30 min at 4°C. The pellet was washed with 200 µl of chilled 70% ethanol, air-dried and resuspended in 30 µl of DEPC-treated milli-Q water. RNA integrity and purity were confirmed on the NanoDrop. Residual genomic DNA was removed by in-solution DNase I digestion. Multiplexed library was generated from 560 ng of fragmented and barcoded RNA as previously described [60].

RNAseq libraries were sequenced on NovSeqX (Illumina, San Diego, USA) in paired-end mode with a total of 300 cycles and sequencing depth of 200 million. The quality of the raw sequencing reads was assessed using FastQC ([61]; version 0.11.9). Trimmomatic ([62]; version 0.39) was used to remove potential adapter reads and perform head-cropping of low-quality bases. Processed reads were mapped to *Methyloversatilis* and PAO1 reference genome (RefSeq accession numbers GCA_059552495.1 and GCF_000006765.1) retrieved from NCBI using bowtie2 ([63] version 2.4.4). FeatureCounts ([64]; version 2.0.3) quantified sequencing reads per gene for genomic feature annotation. Downstream analyses, including normalization, correlation assessment, and identification of differentially expressed genes (false discovery rate, FDR < 0.05 for PAO1; FDR < 0.1 for *Methyloversatilis*), were carried out in R ([52]; version 4.4.3) using the edgeR package ([65]; version 4.0.16) and customized script. Figures were generated in R, and known genes in pyoverdine synthesis and transport in PAO1 were obtained from literature [66].

## Supporting information

Supplementary datafile - Transcriptomics Pseudomonnas PAO1s

Supplementary datafile - Transcriptomics Methyloversatilis 146-33

Supplementary figures and tables with legends

## Acknowledgement

This work was financed by Czech Science Foundation (22-05478S - Iron monopolization versus community service: the two faces of cyanobacterial beta-hydroxy aspartate lipopeptides) – to P.H., Grant from Swiss National Science Foundation (212266) – to R.K., and by the Grant Agency of the University of South Bohemia (GAJU), project no. 126/2024/P, “Do heterotrophic bacteria hijack cyanobacterial siderophores via specific transport mechanisms?” – to B.P.F. Additional support was provided by OP JAK project “Photomachines” Reg. No CZ.02.01.01/00/22_008/0004624 (Czech Ministry of Education, Youth, and Sports (MEYS)), financed by the Ministry of Education, Youth and Sports.

## References

1. Jungblut AD. Ecology and Biogeography of the Cyanobacteria. In: Whitman WB (ed.), Bergey’s Manual of Systematics of Archaea and Bacteria, 1st edn. Wiley, 2022, 1–11.

2. Kranzler C, et al. Iron in Cyanobacteria. Advances in Botanical Research. Elsevier, 2013, 57–105.

3. Jiang H-B et al. New insights into iron acquisition by cyanobacteria: an essential role for ExbB-ExbD complex in inorganic iron uptake. ISME J 2015;9:297–309. 10.1038/ismej.2014.123

4. Burford MA et al. Understanding the relationship between nutrient availability and freshwater cyanobacterial growth and abundance. Inland Waters 2023;13:143–152. 10.1080/20442041.2023.2204050

5. Qiu G-W et al. Iron transport in cyanobacteria – from molecules to communities. Trends Microbiol 2022;30:229–240. 10.1016/j.tim.2021.06.001

6. Rout GR, Sahoo S. ROLE OF IRON IN PLANT GROWTH AND METABOLISM. Rev Agric Sci 2015;3:1–24. 10.7831/ras.3.1

7. Hunnestad AV et al. From the Ocean to the Lab—Assessing Iron Limitation in Cyanobacteria: An Interface Paper. Microorganisms 2020;8:1889. 10.3390/microorganisms8121889

8. Colombo C et al. Review on iron availability in soil: interaction of Fe minerals, plants, and microbes. J Soils Sediments 2014;14:538–548. 10.1007/s11368-013-0814-z

9. Årstøl E, Hohmann-Marriott MF. Cyanobacterial Siderophores—Physiology, Structure, Biosynthesis, and Applications. Mar Drugs 2019;17:281. 10.3390/md17050281

10. Jin Z et al. Conditional privatization of a public siderophore enables Pseudomonas aeruginosa to resist cheater invasion. Nat Commun 2018;9:1383. 10.1038/s41467-018-03791-y

11. Figueiredo ART et al. Siderophores drive invasion dynamics in bacterial communities through their dual role as public good versus public bad. Ecol Lett 2022;25:138–150. 10.1111/ele.13912

12. Kramer J, Özkaya Ö, Kümmerli R. Bacterial siderophores in community and host interactions. Nat Rev Microbiol 2020;18:152–163. 10.1038/s41579-019-0284-4

13. Will V et al. Siderophore specificities of the *Pseudomonas aeruginosa* TonB-dependent transporters ChtA and ActA. FEBS Lett 2023;597:2963–2974. 10.1002/1873-3468.14740

14. Kumar R, Singh A, Srivastava A. Xenosiderophores: bridging the gap in microbial iron acquisition strategies. World J Microbiol Biotechnol 2025;41:69. 10.1007/s11274-025-04287-w

15. Rawat D et al. Iron-dependent mutualism between *Chlorella sorokiniana* and *Ralstonia pickettii* forms the basis for a sustainable bioremediation system. ISME Commun 2022;2:83. 10.1038/s43705-022-00161-0

16. Jiang Y et al. Potential siderophore-dependent mutualism in the harmful dinoflagellate Alexandrium pacificum (Group IV) and bacterium Photobacterium sp. TY1-4 under iron-limited conditions. Harmful Algae 2024;139:102726. 10.1016/j.hal.2024.102726

17. Guan LL, Kanoh K, Kamino K. Effect of Exogenous Siderophores on Iron Uptake Activity of Marine Bacteria under Iron-Limited Conditions. Appl Environ Microbiol 2001;67:1710–1717. 10.1128/AEM.67.4.1710-1717.2001

18. Lucius S, Hagemann M. The primary carbon metabolism in cyanobacteria and its regulation. Front Plant Sci 2024;15:1417680. 10.3389/fpls.2024.1417680

19. Guljamow A et al. High-Density Cultivation of Terrestrial Nostoc Strains Leads to Reprogramming of Secondary Metabolome. Appl Environ Microbiol 2017;83:e01510–17. 10.1128/AEM.01510-17

20. Galica T et al. Cyanochelins, an Overlooked Class of Widely Distributed Cyanobacterial Siderophores, Discovered by Silent Gene Cluster Awakening. Appl Environ Microbiol 2021;87:e03128–20. 10.1128/AEM.03128-20

21. Falcao BP et al. Cyanochelin B: a cyanobacterium-produced siderophore with photolytic properties that negate iron monopolization in UV light. Appl Environ Microbiol 2025;91:e02566–24. 10.1128/aem.02566-24

22. Ringel MT, Brüser T. The biosynthesis of pyoverdines. Microb Cell 2018;5:424–437. 10.15698/mic2018.10.649

23. Balasubramanian D, Mathee K. Comparative transcriptome analyses of Pseudomonas aeruginosa. Hum Genomics 2009;3:349. 10.1186/1479-7364-3-4-361

24. Schalk IJ, Guillon L. Pyoverdine biosynthesis and secretion in *P seudomonas aeruginosa* : implications for metal homeostasis. Environ Microbiol 2013;15:1661–1673. 10.1111/1462-2920.12013

25. Bhamidimarri SP et al. Acquisition of ionic copper by the bacterial outer membrane protein OprC through a novel binding site. PLOS Biol 2021;19:e3001446. 10.1371/journal.pbio.3001446

26. Perraud Q et al. Phenotypic Adaption of Pseudomonas aeruginosa by Hacking Siderophores Produced by Other Microorganisms. Mol Cell Proteomics 2020;19:589–607. 10.1074/mcp.RA119.001829

27. Elias S, Degtyar E, Banin E. FvbA is required for vibriobactin utilization in Pseudomonas aeruginosa. Microbiology 2011;157:2172–2180. 10.1099/mic.0.044768-0

28. Kalyuzhnaya MG et al. Methyloversatilis universalis gen. nov., sp. nov., a novel taxon within the Betaproteobacteria represented by three methylotrophic isolates. Int J Syst Evol Microbiol 2006;56:2517–2522. 10.1099/ijs.0.64422-0

29. Gu S et al. Forging the iron-net: Towards a quantitative understanding of microbial communities via siderophore-mediated interactions. Quant Biol 2025;13:e84. 10.1002/qub2.84

30. He R et al. SIDERITE: Unveiling hidden siderophore diversity in the chemical space through digital exploration. iMeta 2024;3:e192. 10.1002/imt2.192

31. He R et al. Tracing Siderophore Precursors to Primary Metabolism for Ecological Applications. 2025. 10.1101/2025.04.30.651379

32. Masloub JA, Avalon NE. Diversity in Structure and Function: How Cyanobacterial Metallophores Reveal a Broadening Perspective of Microbial Metal-Chelators. J Nat Prod 2026;89:832–845. 10.1021/acs.jnatprod.5c01562

33. Schalk IJ, Perraud Q. *PSEUDOMONAS AERUGINOSA* and its multiple strategies to access iron. Environ Microbiol 2023;25:811–831. 10.1111/1462-2920.16328

34. Chan DCK, Burrows LL. Pseudomonas aeruginosa FpvB Is a High-Affinity Transporter for Xenosiderophores Ferrichrome and Ferrioxamine B. mBio 2023;14:e03149–22. 10.1128/mbio.03149-22

35. Moynié L et al. The complex of ferric-enterobactin with its transporter from Pseudomonas aeruginosa suggests a two-site model. Nat Commun 2019;10:3673. 10.1038/s41467-019-11508-y

36. Miralles I, Ortega R, Montero-Calasanz MDC. Functional and biotechnological potential of microbiome associated with soils colonised by cyanobacteria in drylands. Appl Soil Ecol 2023;192:105076. 10.1016/j.apsoil.2023.105076

37. Stanojković A et al. The global speciation continuum of the cyanobacterium Microcoleus. Nat Commun 2024;15:2122. 10.1038/s41467-024-46459-6

38. Stanojković A et al. Geography and climate drive the distribution and diversification of the cosmopolitan cyanobacterium *Microcoleus* (Oscillatoriales, Cyanobacteria). Eur J Phycol 2022;57:396–405. 10.1080/09670262.2021.2007420

39. microbeAtlas. Available at: https://microbeatlas.org. (accessed on 26 July 2026).

40. Tostado-Islas O et al. Iron limitation by transferrin promotes simultaneous cheating of pyoverdine and exoprotease in *Pseudomonas aeruginosa*. ISME J 2021;15:2379–2389. 10.1038/s41396-021-00938-6

41. D’Onofrio A et al. Siderophores from Neighboring Organisms Promote the Growth of Uncultured Bacteria. Chem Biol 2010;17:254–264. 10.1016/j.chembiol.2010.02.010

42. Rudolph G, Hennecke H, Fischer H-M. Beyond the Fur paradigm: iron-controlled gene expression in rhizobia. FEMS Microbiol Rev 2006;30:631–648. 10.1111/j.1574-6976.2006.00030.x

43. Maza-Márquez P et al. Millimeter-scale vertical partitioning of nitrogen cycling in hypersaline mats reveals prominence of genes encoding multi-heme and prismane proteins. ISME J 2022;16:1119–1129. 10.1038/s41396-021-01161-z

44. Krakau S et al. nf-core/mag: a best-practice pipeline for metagenome hybrid assembly and binning. NAR Genomics Bioinforma 2022;4:lqac007. 10.1093/nargab/lqac007

45. Li D et al. MEGAHIT: an ultra-fast single-node solution for large and complex metagenomics assembly via succinct *de Bruijn* graph. Bioinformatics 2015;31:1674–1676. 10.1093/bioinformatics/btv033

46. Wu Y-W, Simmons BA, Singer SW. MaxBin 2.0: an automated binning algorithm to recover genomes from multiple metagenomic datasets. Bioinformatics 2016;32:605–607. 10.1093/bioinformatics/btv638

47. Chaumeil P-A et al. GTDB-Tk v2: memory friendly classification with the genome taxonomy database. Bioinformatics 2022;38:5315–5316. 10.1093/bioinformatics/btac672

48. Blin K et al. antiSMASH 8.0: extended gene cluster detection capabilities and analyses of chemistry, enzymology, and regulation. Nucleic Acids Res 2025;53:W32–W38. 10.1093/nar/gkaf334

49. Zdouc MM et al. MIBiG 4.0: advancing biosynthetic gene cluster curation through global collaboration. Nucleic Acids Res 2025;53:D678–D690. 10.1093/nar/gkae1115

50. Lu S et al. CDD/SPARCLE: the conserved domain database in 2020. Nucleic Acids Res 2020;48:D265–D268. 10.1093/nar/gkz991

51. Woodcroft BJ et al. Comprehensive taxonomic identification of microbial species in metagenomic data using SingleM and Sandpiper. Nat Biotechnol 2026;44:948– 953. 10.1038/s41587-025-02738-1

52. R Core Team. R: A Language and Environment for Statistical Computing. 2024. Vienna, Austria, 2024.

53. Bankevich A et al. SPAdes: A New Genome Assembly Algorithm and Its Applications to Single-Cell Sequencing. J Comput Biol 2012;19:455–477. 10.1089/cmb.2012.0021

54. Galica T, Hrouzek P, Mareš J. Genome mining reveals high incidence of putative lipopeptide biosynthesis NRPS / PKS clusters containing fatty acyl-AMP ligase genes in biofilm-forming cyanobacteria. J Phycol 2017;53:985–998. 10.1111/jpy.12555

55. Gilchrist CLM, Chooi Y-H. clinker & clustermap.js: automatic generation of gene cluster comparison figures. Bioinformatics 2021;37:2473–2475. 10.1093/bioinformatics/btab007

56. Wickham H. ggplot2: elegant graphics for data analysis, Second edition. Cham: Springer international publishing, 2016.

57. Koblitz J. 1341: MIN E - METHYLOVERSATILIS MEDIUM | Media | MediaDive. https://mediadive.dsmz.de. .

58. Pinto FL et al. Analysis of current and alternative phenol based RNA extraction methodologies for cyanobacteria. BMC Mol Biol 2009;10:79. 10.1186/1471-2199-10-79

59. Krynická V et al. Depletion of the FtsH1/3 Proteolytic Complex Suppresses the Nutrient Stress Response in the Cyanobacterium *Synechocystis* sp strain PCC 6803. Plant Cell 2019;31:2912–2928. 10.1105/tpc.19.00411

60. Shishkin AA et al. Simultaneous generation of many RNA-seq libraries in a single reaction. Nat Methods 2015;12:323–325. 10.1038/nmeth.3313

61. Babraham Bioinformatics - FastQC A Quality Control tool for High Throughput Sequence Data. https://www.bioinformatics.babraham.ac.uk/projects/fastqc/. .

62. Bolger AM, Lohse M, Usadel B. Trimmomatic: a flexible trimmer for Illumina sequence data. Bioinformatics 2014;30:2114–2120. 10.1093/bioinformatics/btu170

63. Langmead B, Salzberg SL. Fast gapped-read alignment with Bowtie 2. Nat Methods 2012;9:357–359. 10.1038/nmeth.1923

64. Liao Y, Smyth GK, Shi W. featureCounts: an efficient general purpose program for assigning sequence reads to genomic features. Bioinformatics 2014;30:923– 930. 10.1093/bioinformatics/btt656

65. hen Y, et al. edgeR v4: powerful differential analysis of sequencing data with expanded functionality and improved support for small counts and larger datasets. Nucleic Acids Res 2025;53:gkaf018. 10.1093/nar/gkaf018

66. Lamont IL, Martin LW. Identification and characterization of novel pyoverdine synthesis genes in Pseudomonas aeruginosa. Microbiology 2003;149:833–842. 10.1099/mic.0.26085-0

