## Supplementary figures and tables with legends for "Utilization of cyanobacterial siderophore cyanochelin B by phylogenetically distant heterotrophs suggest its role in mediating microbial interactions"

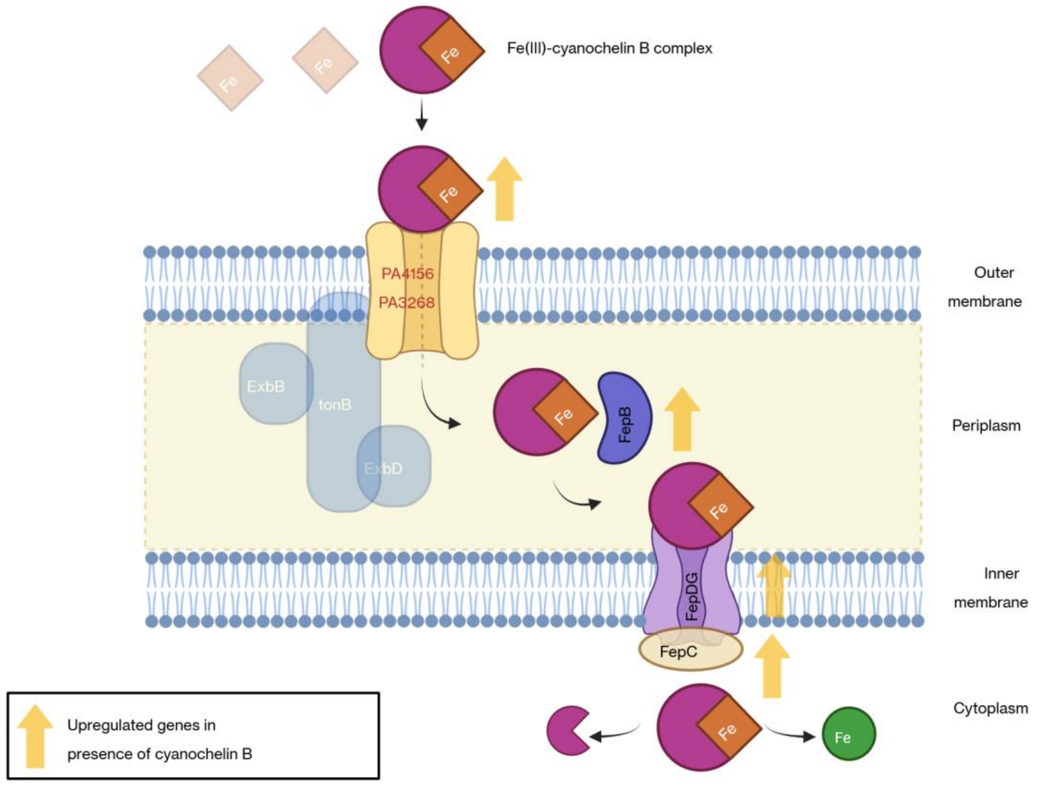

***Figure S1***: Hypothetical model for internalization of cyanochelin B during iron starvation by PAO1 strain. The upward facing arrows in the schematic represent genes upregulated in presence of CychB.

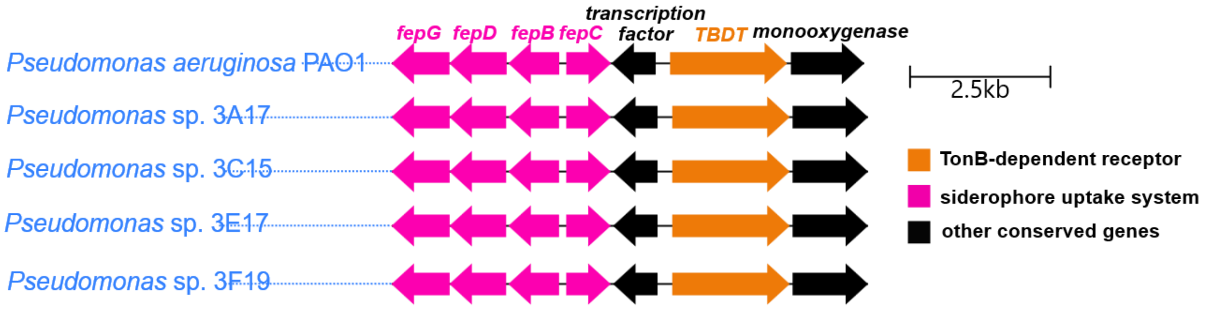

***Figure S2***: Clinker-based comparison of the target iron gene clusters of the hypothetical gene cluster shown in figure S1 found in PAO1. Homologous genes in *pseudomonas* isolates are color coded and gene orientation is indicated by direction of the arrow direction.

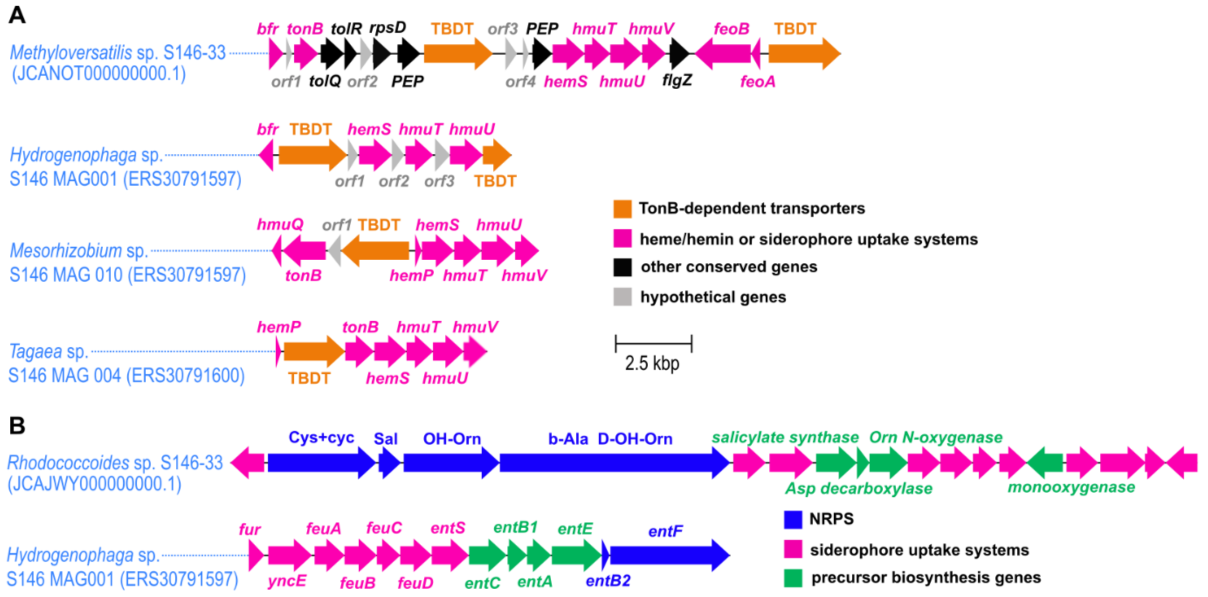

***Figure S3***: Genetic potential for TonB-dependent siderophore uptake and siderophore synthesis in bacteria from sample S146. **A,** Putative TonB-dependent siderophore uptake gene cassettes in the *Methyloversatilis* sp. S146-33 strain isolated from sample S146 compared to similar gene cassettes identified in cohabitant MAGs from the *Phormidesmis* sp. S146-33 culture. **B,** Putative siderophore biosynthetic gene clusters identified in strain *Rhodococcoides* sp, S146-33 and *Hydrogenophaga* sp. MAG. The metallophore BGC in the *Rhodococcoides* genome contains a set of genes typical for mycobactin/madurastatin-type hydroxamate siderophores, including the genes for biosynthesis of salicylate and beta-alanine precursors, the NRPS machinery for peptide backbone assembly, and associated siderophore transport/uptake. The *Hydrogenophaga* genome contains a set of entA,B,C,E, and entF genes required for enterobactin/catecholate biosynthesis as well as the associated uptake/transport machinery; entD (phosphopanteteine transferase) homolog was identified in a different contig within the *Hydrogenophaga* MAG. NRPS - non-ribosomal peptide synthase, TBDR - TonB-dependent receptor.

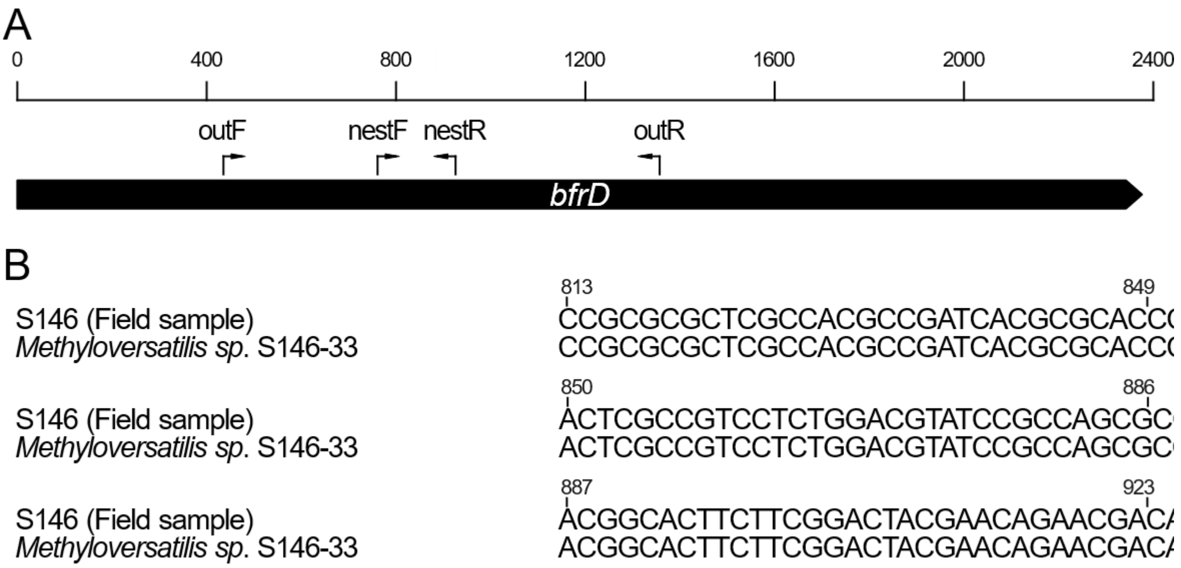

***Figure S4***: A, Map of the annealing sites of the nested PCR primers designed for bfrD gene, a putative TonB-dependent receptor encoded in *Methyloversatilis* sp. 146-33 genome. B, Alignment of the nested amplicon produced from the field sample 146 and its target sequence in the *Methyloversatilis* sp. 146-33 genome, both sequences are identical.

### SUPPLEMENTARY TABLES

| Sr no | ID | Source | Pyoverdine production |
|---|---|---|---|
| 1 | **3F19** | Pond | + |
| 2 | **3E17** | Pond | + |
| 3 | **3A17** | Pond | + |
| 4 | **3C15** | Pond | + |
| 5 | **3A10** | Pond | + |
| 6 | **s3e11** | Soil | - |
| 7 | **3H02** | Pond | - |
| 8 | **s3b15** | Soil | - |
| 9 | **s3b19** | Soil | - |
| 10 | **s3b09** | Soil | + |
| 11 | **3A19** | Pond | + |
| 12 | **3D20** | Pond | + |
| 13 | **3G15** | Pond | + |
| 14 | **s3d6** | Soil | + |
| 15 | **3B17** | Pond | + |

***Supplementary Table 1.*** Natural isolates of Pseudomonas isolated from Irchel campus in University of Zurich and used in this study, including species ID, source of isolation, and pyoverdine production profile.

| COMPONENTS | M (G/L) | COMMENTS |
|---|---|---|
| NANO3 | 46.7 | |
| CA(NO3)2.4H2O | 5.9 | |
| K2HPO4 | 3.1 | Add after autoclaving! |
| MGSO4.7H2O | 2.5 | |
| NA2CO3 | 2.1 | |
| FE-EDTA SOLUTION | | |
| AFTER ADDING THE MAIN COMPONENTS ADD 0.08 ML OF TRACE ELEMENT SOLUTION TO 1 LITRE MEDIUM. INDIVIDUAL COMPONENTS ARE AS FOLLOWS: | | |
| TRACE ELEMENTS | mg/L | |
| H3BO3 | 0.09008 | |
| MNSO4.4H2O | 0.075065 | |
| NA2WO4.2H2O | 0.05502 | |
| (NH4)6MO7O24.4H2O | 0.14705 | |
| KBR | 0.066035 | |
| KI | 0.06804 | |
| ZNSO4.7H2O | 0.135075 | |
| CD(NO3)2.4H2O | 0.022121 | |
| CO(NO3)2.6H2O | 0.0460237 | |
| CUSO4.5H2O | 0.0303 | |
| NISO4(NH4)2SO4.6H2O | 0.06257 | |
| CR(NO3)2.7H2O | 0.023035 | |
| V2O4(SO4)3.16H2O | 0.079055 | |
| AL2(SO4)3K2SO4.24 H2O | 0.09008 | |
| C & N MIX | mg/L | |
| D-GLUCOSE | 0.09008 | |
| D-RIBOSE | 0.075065 | |
| SODIUM PYRUVATE | 0.05502 | |
| SODIUM CITRATE | 0.14705 | |
| OXALOACETIC ACID | 0.066035 | |
| SODIUM ACETATE | 0.06804 | |
| SODIUM SUCCINATE | 0.135075 | |
| N-ACETYLGLUCOSAMINE | 0.022121 | |
| GLYCEROL | 0.0460237 | |
| UREA | 0.0303 | |
| TAURINE | 0.06257 | |
| ETHANOL | 0.023035 | |
| SODIUM THIOSULFATE | 0.079055 | |
| AMINO ACID MIX | µg/L | |
| ISOLEUCINE | 26.2347 | |
| LEUCINE | 26.2347 | |
| LYSINE | 29.2376 | |
| METHIONINE | 29.8425 | |
| PHENYLALANINE | 33.038 | |

| | |
|---|---|
| **THREONINE** | 23.8239 |
| **TRYPTOPHAN** | 40.8452 |
| **VALINE** | 23.4294 |
| **ARGININE** | 34.8403 |
| **HISTIDINE** | 31.031 |
| **ALANINE** | 17.8187 |
| **ASPARAGINE** | 26.4237 |
| **ASPARTATE** | 26.6206 |
| **CYSTEINE** | 24.2318 |
| **GLUTAMATE** | 58.852 |
| **GLUTAMINE** | 58.458 |
| **GLYCINE** | 15.0134 |
| **PROLINE** | 23.0262 |
| **SERINE** | 21.0186 |
| **TYROSINE** | 36.2379 |
| **VITAMIN MIX** | **µg/L** |
| **V B1 THIAMINE** | 200.001 |
| **V B3 NIACIN** | 9.7696 |
| **V B12 COBALAMINE** | 0.1003 |
| **V BX PARA-AMINO BENZOIC ACID** | 0.6857 |
| **V B6 PYRIDOXINE** | 15.2174 |
| **V B5 PANTOTHENIC ACID** | 19.5372 |
| **V B7 BIOTIN** | 0.97724 |
| **V B9 FOLSÄURE** | 1.7656 |
| **MYO-INOSITOL** | 99.9888 |
| **V B2 RIBOFLAVIN** | 3.7636 |

***Supplementary Table 2.*** Media recipe for Z-modified medium used for cohabitant bacteria isolation from *Phormidesmis* enriched culture.

### SUPPLEMENTARY DATAFILES

**Supplementary datafile 1.** Gene-level transcriptomic data for *Pseudomonas* (PAO1) after feeding with cyanochelin B (and control) highlighting genomic annotation, locus tag and transcriptomic profile ($log_2$ fold-changes for all conditions, FDR-adjusted significance values, and expression counts for individual biological replicates).

**Supplementary datafile 2.** Gene-level transcriptomic data for *Methyloversatilis* sp. S146-33 after feeding with cyanochelin B (and control) highlighting genomic annotation, locus tag and transcriptomic profile ($log_2$ fold-changes for all conditions, FDR-adjusted significance values, and expression counts for individual biological replicates).
